# Combination of Cas9 and adeno-associated vectors (AAVs) enables efficient *in vivo* knockdown of precise miRNAs in the rodent brain

**DOI:** 10.1101/2025.03.26.645225

**Authors:** David Roura-Martinez, Natalia Popa, Florence Jaouen, Cynthia Rombaut, Catherine Lepolard, Dipankar Bachar, Ana Borges, Maxime Cazorla, Maxime Villet, Sebastien Moreno, Hélène Marie, Eduardo Gascon

## Abstract

Although the advent of Cas9 technology has expanded our ability to precisely edit the genome, manipulating microRNAs *in vivo* has been shown to be particularly challenging, especially in the brain. Here, we sought to generate novel tools aiming at targeting and efficiently downregulating defined microRNAs species in a cell-specific manner so that their function in discrete neuronal networks could be investigated. Focusing on miR-124, a microRNA highly expressed in the mammalian brain and transcribed from three independent chromosomal loci, we designed and validated different guide RNAs directed against this miRNA. *In vitro*, our Cas9 designs show not only a significant reduction in miR-124 levels but also a functional effect on miR-124 silencing. Similarly, when packed into AAV vectors and injected into the mouse cortex, miR-124-Cas9 vectors strongly downregulate miR-124 levels without affecting the expression of other miRNAs. In parallel, levels of endogenous miR-124 targets exhibit a significant increase supporting the release of its silencing activity. To functionally validate our tools, we provide evidences that deletion of miR-124 in the subventricular zone altered migration of newly generated neurons into the olfactory bulb. Finally, we also showed that our vectors modified the Ca^2+^ permeability of AMPA receptors, a robust functional output downstream of miR-124. These tools are expected to help elucidating miRNA function in complex experimental settings such as brain networks *in vivo*.

## Introduction

microRNAs (miRNAs) are a class of short (20-25 nt) non-coding RNAs (Izaurralde, 2015; Huntzinger and Izaurralde, 2011) silencing gene expression at the post-transcriptional level. miRNAs have been involved in multiple biological processes (Papagiannakopoulos and Kosik, 2008; Kosik, 2006; Ambros, 2011)as well as in a variety of pathological conditions (Visone and Croce, 2009; Abu-Elneel et al., 2008). Although accumulating evidence points out a major role of miRNAs in the brain (Salta and De Strooper, 2017; Issler and Chen, 2015), our understanding of miRNAs functions remains scarce.

Loss-of-function studies are a fundamental strategy to investigate the function of coding genes as well as of miRNAs. However, the lack of tools to effectively delete miRNAs in specific networks has precluded careful interrogation of miRNA function in the brain (Baigude and Rana, 2014; Stenvang et al., 2012). On one hand, strategies based on miRNA inhibition (e.g. antagomirs or sponges) effectively repress miRNA activity *in vivo* (Elmén et al., 2008; Krützfeldt et al., 2005) but are difficult to target to specific cell type/circuits. On the other hand, classical genetics, extensively used to downregulate/upregulate levels of coding genes, have enormous limitations when applied to miRNAs. Indeed, owing to chromosomal arrangement (e.g. miRNAs embedded within a coding gene or in miRNA clusters) and transcription from multiple genomic loci (e.g. three miR-124 genes exist in mouse and human genome), very few miRNAs knockout mouse models have been successfully generated in the last decade (Sanuki et al., 2011; Tan et al., 2013; Liu et al., 2022). In that context, the advent of genome editing technologies based on bacterial Cas9 nuclease has significantly enhanced our ability to carry out targeted genetic manipulations. In addition, Cas9 possesses an intrinsic multiplex activity and is able to target the same sequence if found at different chromosomal locations (Niu et al., 2017)

Here, we developed guide RNAs (gRNAs) designs to inactivate miR-124, a conserved miRNA highly expressed in the mammalian nervous system (Gao, 2010), playing key functions during development (Rajman and Schratt, 2017) and involved in several pathological processes (Roy et al., 2017; Gascon et al., 2014). We systematically investigated their efficiency *in vitro* using different cell lines engineered to express miR-124 from different chromosomal loci. Then, we tested our tools in different *in vivo* settings. Using adeno-associated viral vectors (AAVs) as delivery tool, we show that our Cas9 designs can disrupt miR-124 expression in the cortex of adult mice and alter the levels of downstream target genes. Moreover, as previously reported, we observed that Cas9-mediated downregulation of miR-124: i) affected the Ca^2+^-permeability of AMPA receptors (AMPARs) in hippocampal neurons; and ii) altered migration of newly generated neurons in the olfactory bulb. Together, these observations provide a functional validation of our tools and opens new avenues for investigating microRNA function in the brain via targeted inactivation in precise neuronal networks.

## Results

### gRNA designs to inactivate miR-124 in vitro

Several previous reports have shown that Cas9-induced insertion-deletions (indels) result in miRNA inactivation (Ho et al., 2015). We therefore designed four different gRNAs targeting mouse miR-124 and tested their efficiency *in vitro*. First, we transfected our different gRNAs together with Cas9 in HEK cells engineered to stably express mouse miR-124-1 or 2 (HEK^m124-1^ and HEK^m124-2^) (Lepolard et al., 2023). Compared to the naïve cells or those transfected with a plasmid driving Cas9 but no gRNA, our designs show a variable degree of miR-124 downregulation (Suppl. Fig 1A) depending on miR-124 isoform. Interestingly, one of our designs (gRNA1) showed no activity whereas another (gRNA4) exhibited a similar efficiency in targeting both miR-124-1 and 2. Both gRNAs were selected for further analysis.

miRNAs silence gene expression post-transcriptionally so that reduced levels of a miRNA result in a parallel increase of its downstream targets. To confirm the functional impact of miR-124 downregulation observed with our Cas9 approach, we constructed HEK lines co-expressing both miR-124 and a miR-124 reporter (an RFP reporter bearing miR-124 target sequences, HEK^m124-1-RFP^ and HEK^m124-2-RFP^) (Lepolard et al., 2023). If functionally relevant, Cas9 targeting miR-124 in these cell lines should lead to an increase of reporter fluorescence. We therefore transfected HEK^124-1-RFP^ and HEK^124-2-RFP^ with plasmids driving simultaneously the expression of Cas9, a control CFP reporter and either gRNA1 or gRNA4 and analyzed fluorescence at different time points. CFP was used to identify transfected Cas9^+^ cells and also as a control reporter, not submitted to miR-124 regulation. (Suppl Fig1B). We observed that RFP in cells transfected with gRNA1 (CFP^+^) was not different compared to those untransfected (CFP^-^) at any time point. In contrast, cells having received gRNA4 exhibited a temporal increase in reporter fluorescence in CFP^+^ cells, being significant from 10 days after transfection (Suppl Fig 1C). These results confirmed that the observed reduction in miR-124 levels after Cas9 deletion are functionally relevant at the protein level (Huang et al., 2023).

### Cas9-miR-124 constructs packed into AAVs inactivate miR-124 in the mouse cortex

To test the ability of our designs to inactivate miR-124 in the brain, we generated adeno-associated vectors (AAVs) driving the concomitant expression of the saCas9 and the gRNA as previously described (Ran et al., 2015). We produced an AAV containing the efficient gRNA (gRNA#4, AAV-Cas9-miR-124^g4^) and two control vectors, one driving the inefficient gRNA (gRNA#1, AAV-Cas9-miR-124^g1^) and another which does not express any gRNA (AAV-Cas9-miR-124^empty^) (Fig 1A). Those vectors were injected into the motor cortex of wild-type mice. Four to six weeks later, animals were sacrificed and used for different downstream applications including DNA sequencing, in situ hybridization, miRNA quantification and RNA sequencing. A summary of the different experimental designs can be found in Suppl Fig 2.

**Figure 1.**
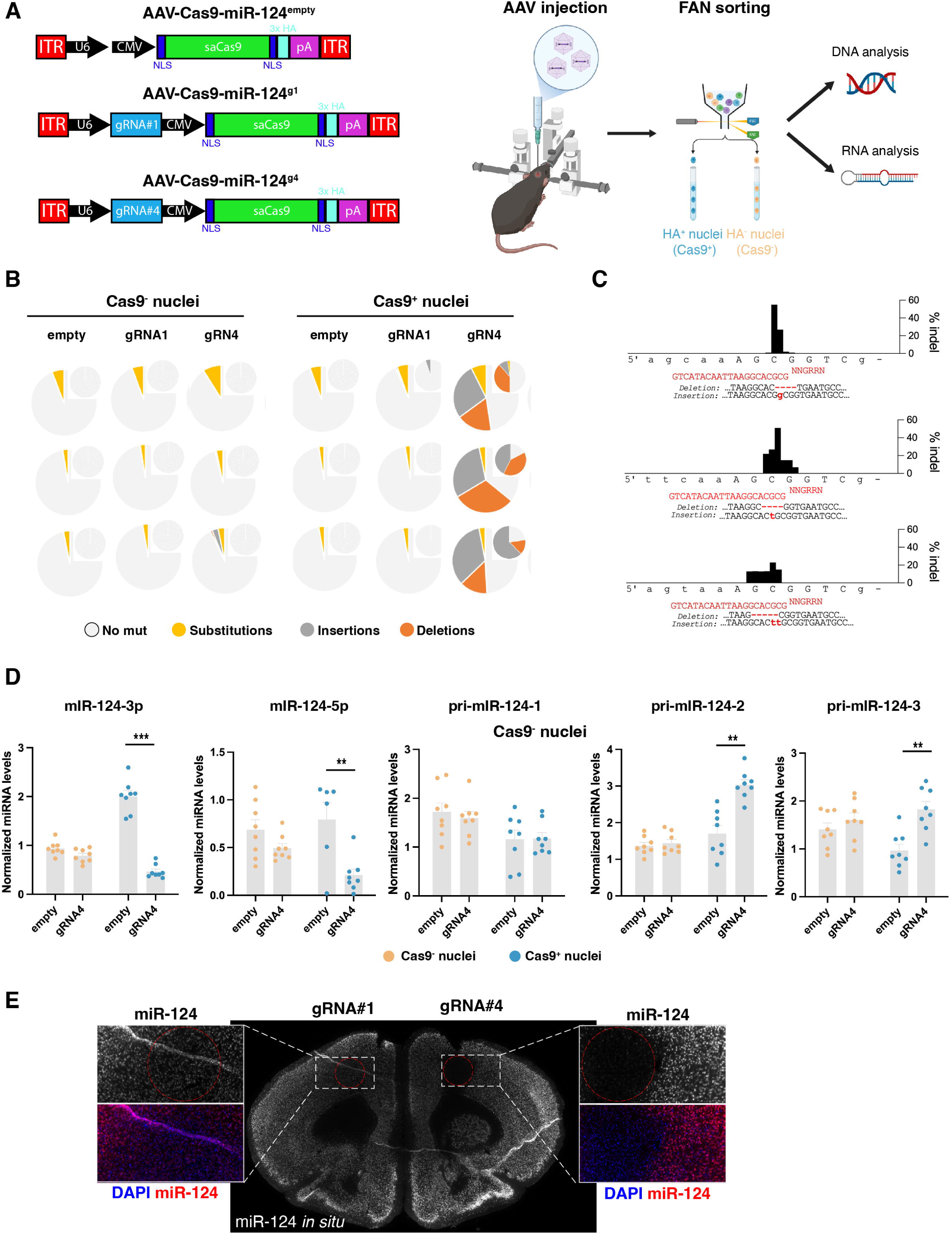
Cas9-mediated inactivation of miR-124 in the mouse brain A. Schematic representation of the AAVs design used in this study (left) as well as the experimental design for *in vivo* experiments (right). Mice receive a control AAV (empty or g1) on one motor cortex and the g4 on the contralateral side. After 4-6 weeks, most animals were sacrificed and processed for nuclear isolation, FANS sorting and DNA or RNA analysis. Some were transcardially perfused for *in situ* hybridization. B. Cas9-driven double-strands breaks results in mutations at miR-124 loci. Following the experimental approach depicted in panel A, animals having received the gRNA4 show a high proportion (40-70%) of insertion or deletions in all three miR-124 loci. DNA sequencing data are displayed as the main pie charts whereas those obtained from PCR cloning are shown as insert pie chart. Indels were virtually absent in mice injected with control AAVs or in Cas9^-^ nuclei. Substitutions are equally common across all conditions in DNA sequencing data but not in cloning, likely representing technical noise (from multiple cycles of PCR amplification). C. Illustration of the location and proportion of indels induced by gRNA4 at the three miR-124 loci. D. gRNA4 significantly decreases miR-124 levels *in vivo*. Using the same experimental strategy, mi-124 levels were measured using qPCR. As illustrated, a robust downregulation in both miR-124-3p (n=8, 2-way ANOVA, F (1, 14) = 140.5, p<0.0001; Bonferroni test, gRNA4 vs empty p<0.0001) and 5p (n=8, 2-way ANOVA, F (1, 26) = 14.82, p<0.0007; Bonferroni test, gRNA4 vs empty p<0.0011) is observed in Cas9^+^ nuclei for miR-124. No differences in miR-124 levels can be found in Cas9^-^ nuclei. A significant increase of the primary transcript from loci 2 and 3 was also observed in Cas9^+^ nuclei (n=8, 2-way ANOVA, for miR-124-1, F (1, 14) = 0.087, p<0.7715; for miR-124-2, F (1, 14) = 26.55, p=0.0001; Bonferroni test, gRNA4 vs empty p<0.0001Bonferroni test; for miR-124-3, F (1, 14) = 19.5, p=0.0006; Bonferroni test, gRNA4 vs empty p=0.0004) E. Picture of miR-124 *in situ* hybridization in the brain. In an animal having received gRNA1 (left hemisphere) and gRNA4 (right hemisphere), miR-124 signal shows a clear decrease in the area injected with gRNA4 (red circle). Scale bar 50 μm.

We first analyzed the effect of our Cas9 tools at the DNA level. Mice having received the different AAVs were sacrificed and the injected area dissected. Nuclei from this region were extracted and two fractions (Cas9^-^ and Cas9^+^) were isolated via Fluorescence Activated Nuclei Sorting (FANS) (Fig 1A, see methods). We then performed DNA sequencing to quantify mutations in the different miR-124 loci as well as in the top 10 off-targets. We first assessed the ability of our designs (AAV-Cas9-miR-124^g4^, AAV-Cas9-miR-124^g1^ and AAV-Cas9-miR-124^empty^) to target the miR-124 loci. As shown in Fig 1B, AAV-Cas9-miR-124^g4^ induced multiple indels at the expected location (3 nt upstream of the Protospacer Adjacent Motif, PAM, sequence) in all three miR-124 loci. In contrast, AAV-Cas9-miR-124^g1^ and AAV-Cas9-miR- 124^empty^ resulted in very few mutations that might likely represent experimental noise from PCR amplifications and/or sequencing errors. To obtain further confirmation, we amplified by PCR a fragment of 300 nt around the PAM sequence for the three miR- 124 loci and cloned those fragments into a sequencing plasmid. We then sequenced 200 independent clones and for each locus and condition. Our results (Fig. 1B-C) confirmed that only mice injected with the AAV-Cas9-miR-124^g4^ resulted in a high proportion of indels whereas the control AAVs showed no mutations. We then searched for off-target effects of our gRNAs. As expected, owing to limited homology of our gRNA designs (4 mismatches) to other genomic regions, we did not find any indels in the selected off-target loci confirming the specificity of Cas9 editing (Suppl Fig 3).

We next sought to analyze miR-124 expression using quantitative PCR. For that, RNA from FANS-sorted nuclei was extracted. We were able to demonstrate a clear reduction of miR-124-3p and 5p levels in Cas9^+^ nuclei infected with the AAV-Cas9-miR-124^g4^ (Fig. 1D). Importantly, no such effect could be observed in Cas^-^ nuclei. To confirm the specificity of such observations, we ran additional controls. On one hand, neither AAV-Cas9-miR-124^g1^ nor AAV-Cas9-miR-124^empty^ altered miR-124 expression neither in Cas9^-^ or Cas9^+^ cells (Fig 1D and Suppl. Fig 4A). On the other hand, for the efficient guide, the reduction of miRNA levels was restricted to miR-124 but no other miRNAs were affected (Suppl. Fig 4B). Finally, we found that reduction of mature miR-124 was concomitant with an increase in the primary miR-124 transcript (pri-miR-124) levels from both locus 2 and 3 (Fig 1D).

To obtain additional confirmation of the miR-124 targeting, we carried out in situ hybridization on mice having received an injection of either AAV-Cas9-miR-124^g1^ or AAV-Cas9-miR-124^g4^ in the motor cortex. As shown in Fig 1E, a clear downregulation of miR-124 signal was observed in the area injected with the efficient guide but not in the contralateral hemisphere. Together, these data together indicate that AAVs driving Cas9 can target miR-124 in the mouse brain with high efficacy and specificity.

### miR-124 inactivation in vivo alters the expression of downstream targets

miR-124 deletion should result in the deregulation of its downstream targets. To address this issue, we performed RNA sequencing on FANS sorted Cas9^+^ and Cas9^-^nuclei. Using these transcriptomic profiles, we first carried out a hierarchical clustering (Fig 2A). As expected, tissues expressing either AAV clearly showed a different mRNA expression pattern and segregated into two groups.

**Figure 2.**
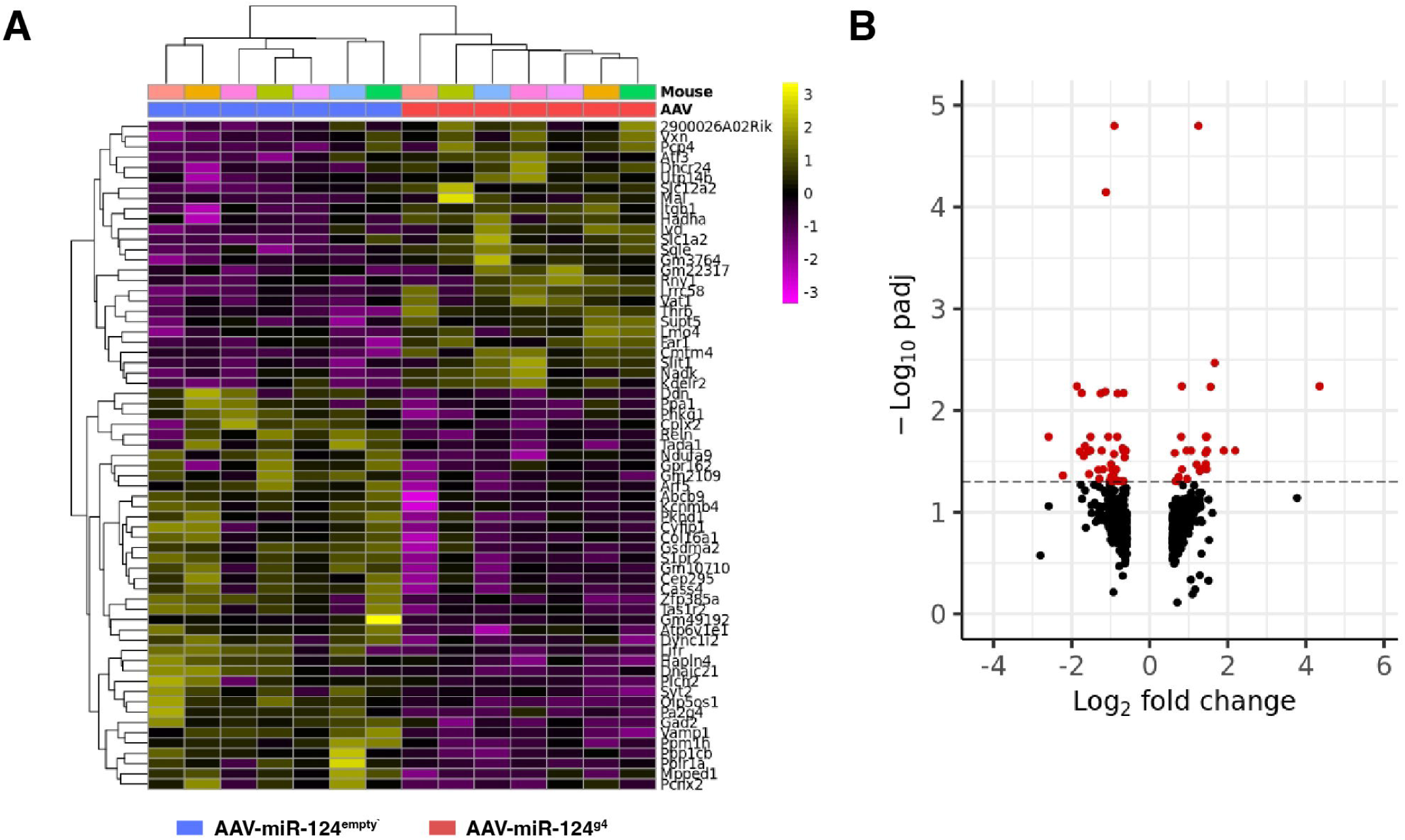
Cas9-mediated inactivation of miR-124 *in vivo* results in deregulation of transcriptomic profiles. A. Hierarchical clustering of mRNA sequencing data obtained from animals injected with AAV-Cas9-miR-124^empty^ and AAV-Cas9-miR-124^g4^ on each motor cortex. As illustrated, in Cas9^+^ nuclei, transcriptomic profiles nicely segregated both groups. This figure was obtained with the pHeatmap function in Rstudio from the count matrix of the 65 most DEGs after regularized logarithm transformation of the data. B. Volcano plots of deregulated transcripts after miR-124 loss. When compared, Cas9^+^ nuclei having received gRNA4 show a significant upregulation of 24 mRNAs whereas 36 are downregulated (DESeq2, FDR, p adjusted< 0.05). The plot was generated with the EnhancedVolcano package in Rstudio from the 660 genes with the highest FC values.

We then analyzed quantitatively these differences between control and gRNA4 cortices. We found that there was almost no transcript differentially expressed among Cas9^-^ nuclei (2 upregulated mRNAs) (Suppl. Fig 5). In contrast, there was a distinct profile of mRNA expression across the Cas9^+^ nuclei; a total of 60 transcripts showed significantly different levels between control and gRNA4-injected cortices (Fig 2B). Since miRNAs silence gene expression through mRNA degradation and/or translation inhibition, we focused on those mRNAs which showed increased levels in gRNA4-injected cortices. Using a well-established tool designed to search for downstream targets (targetScan8) (McGeary et al., 2019), we assessed whether the upregulated transcripts were predicted miR-124 targets. We found that, among 24 upregulated mRNAs, only 5 did not contain miR-124 binding sequences indicating a clear enrichment for miR-124 targets (Table 1). In contrast, only 5 of the 36 downregulated transcripts were found to be miR-124 target arguing for the specificity of Cas9 inactivation on releasing post-transcriptional silencing.

**Table 1.** List of upregulated transcripts in mRNA sequencing experiments and the presence of miR-124 binding sequences.

|  | log2FoldChange | pvalue | padj | miR-124 target |
| --- | --- | --- | --- | --- |
| Lrrc58 | 1,25044498 | 4,16E-08 | 1,59E-05 | Yes |
| Vat1 | 1,66584492 | 2,03E-05 | 0,00340739 | Yes |
| Mal | 4,35188139 | 4,31E-05 | 0,00577354 | No |
| Thrb | 0,8223197 | 5,38E-05 | 0,00577354 | Yes |
| Slit1 | 1,55668644 | 6,97E-05 | 0,00585219 | Yes |
| Itgb1 | 1,44036321 | 0,00043102 | 0,01810288 | Yes |
| Rny1 | 1,46329948 | 0,00053064 | 0,0181029 | Yes |
| Cmtm4 | 0,80849186 | 0,00052396 | 0,0181029 | Yes |
| Kdelr2 | 2,19131924 | 0,00096706 | 0,02481876 | Yes |
| Slc12a2 | 1,90046387 | 0,00099718 | 0,02481876 | Yes |
| 2900026A02Rik | 1,46570632 | 0,00087093 | 0,02481876 | Yes |
| Pcp4 | 0,94513926 | 0,00110744 | 0,02483638 | No |
| Slc1a2 | 0,64111364 | 0,00132587 | 0,02620534 | Yes |
| Utp14b | 1,42942728 | 0,00136724 | 0,02625096 | Yes |
| Sqle | 1,20384829 | 0,00206775 | 0,03389094 | No |
| Atl3 | 1,456768 | 0,00270245 | 0,03783429 | Yes |
| Ivd | 1,30937991 | 0,00269815 | 0,03783429 | No |
| Hadha | 1,30207828 | 0,00264855 | 0,03783429 | Yes |
| Dhcr24 | 0,82600948 | 0,00258223 | 0,03783429 | Yes |
| Nadk | 1,28338698 | 0,00294281 | 0,0395513 | No |
| Lmo4 | 0,74273224 | 0,00368395 | 0,04501115 | Yes |
| Supt5 | 0,95680872 | 0,00398391 | 0,04696824 | Not in database |
| Vxn | 0,72203523 | 0,00442019 | 0,04939675 | Not in database |
| Far1 | 0,66067667 | 0,00477796 | 0,04939675 | Yes |

**Table 1.**
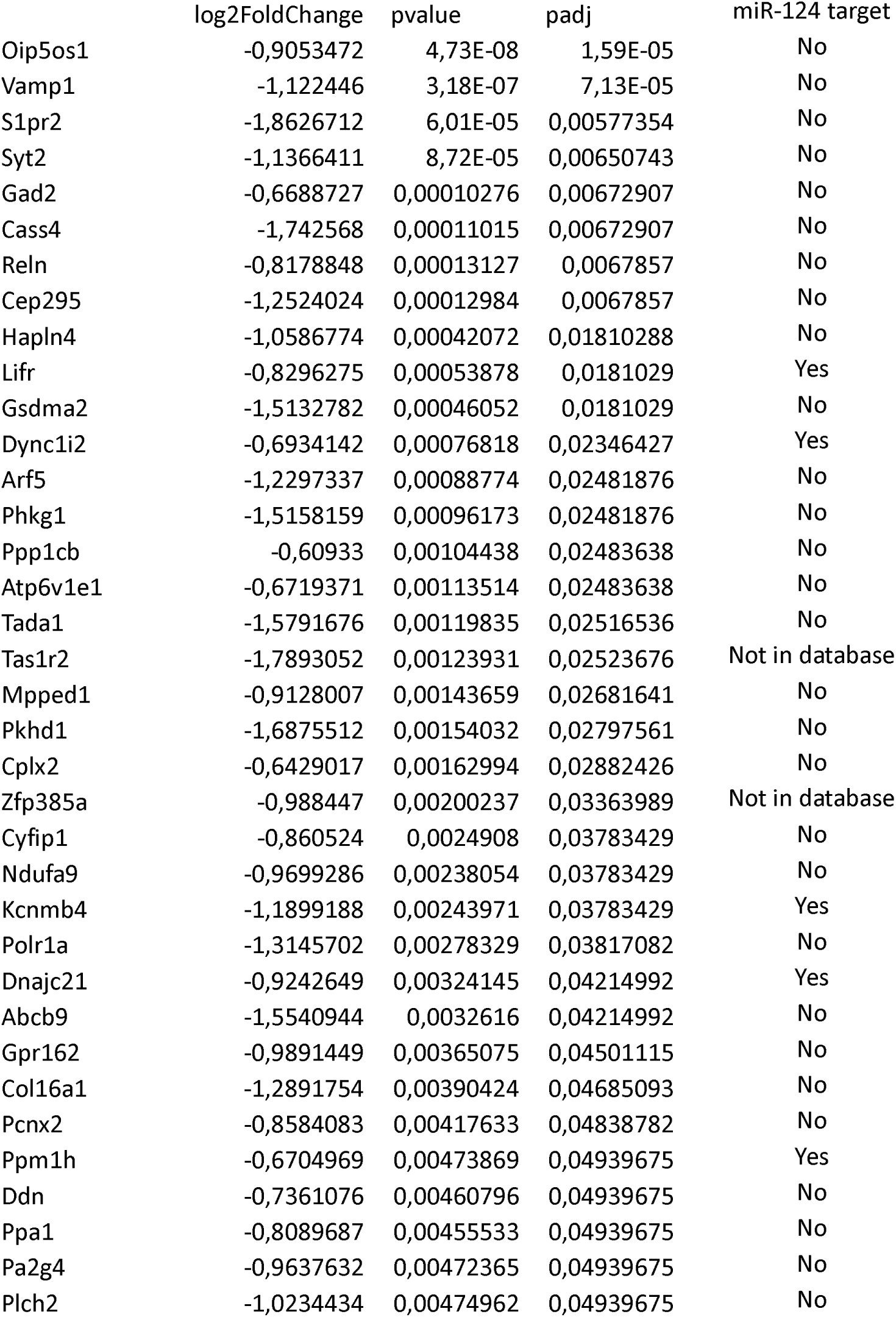
List of downregulated transcripts in mRNA sequencing experiments and the presence of miR-124 binding sequences.

### Functional consequences of miR-124 inactivation in the mouse brain

We next sought to provide a functional proof of principle of miR-124 inactivation in vivo. A region where miR-124 has shown to be essential is the neurogenic niche of the subventricular zone. Previous reports have suggested that treatment with miR- 124 antisense oligonucleotides disrupts migration of neuroblasts generated in this zone and the supply of new neurons *in vivo* to the olfactory bulb (OB) (Cheng et al., 2009). Reporter mice has documented that miR-124 is not expressed neither in slowly dividing neural stem cells (NSCs) nor in rapidly-dividing progenitors (C cells) (Åkerblom et al., 2012). It is well-established that NSC are resident cells in contact with the lateral ventricle so that AAV delivered intraventricularly might enable the genetic deletion of miR-124 in these cells and examine the downstream functional effects on neuroblasts migration in a spatially different brain region. To manipulate miR-124 expression in the SVZ-OB neurogenic niche, we injected either AAV-Cas9-miR-124^g4^ or AAV-Cas9-miR-124^g1^ in the anterior horn of the right lateral ventricle of newborn mice so that a fraction of resident NSCs could be genetically modified. Ten days later, we injected BrdU intraperitoneally to label actively dividing cells. Finally, we analyzed the number of migrating neuroblasts that reached the OB at P21 using immunostaining for BrdU and doublecortin (DCX), a protein known to label immature neurons in the brain (Fig 3A). To have an internal control in these experiments, we quantified the number of BrdU^+^ cells in both olfactory bulbs and calculated the ratio between the right (injected) and left (non-injected) sides. A ratio near 1 suggest no alteration in migration whereas a ratio below 1 is an indication of disrupted migration. The results (Fig 3B) confirm that animals injected with AAV-Cas9-miR-124^g1^ show no differences in migration between left and right hemispheres (ratio 1,14±0.3 n=4/group). In sharp contrast, the group of animals injected with AAV-Cas9-miR- 124^g4^ show a dramatic reduction of the number of newly generated neurons in the injected side but no difference in the contralateral hemisphere. Consequently, the ratio right/left was significantly decreased (0.62±0.13, n=3, p=0.0452). We thus demonstrate that deletion of miR-124 in the SVZ progenitors altered neuroblast migration, in agreement with previous reports using miR-124 antisense (Cheng et al., 2009) and similar to other models in which neuroblast migration is known to be perturbed (Cremer et al., 1994; Saghatelyan et al., 2004).

**Figure 3.**
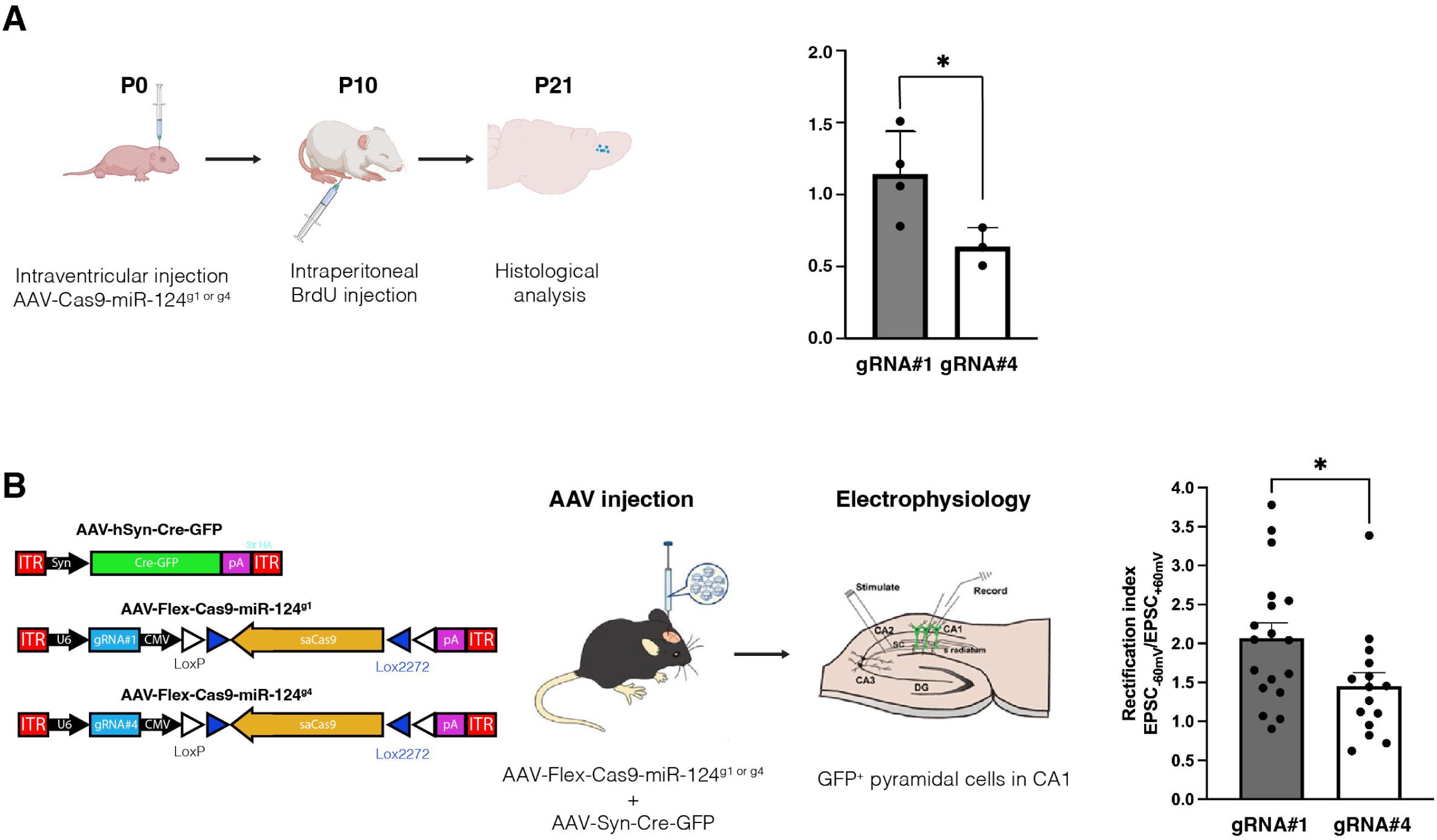
Functional validation of miR-124 loss *in vivo*. A. miR-124 inactivation alters migration of newly generated neurons from the subventricular zone to the olfactory bulb. Schematic representation of the experimental protocol used in these experiments (left). Newborn mice receive first a unilateral injection of AAV-Cas9-miR-124^g1^ or AAV-Cas9-miR-124^g4^ into the lateral ventricle. After 10 days, proliferating cells were labeled using four intraperitoneal injections of BrdU. Migration was evaluated by counting the number of BrdU+ nuclei in the migratory stream at P21. For quantification, the ratio between the injected and non-injected sides for each animal was calculated. As shown, less neuroblasts were found in animals injected with gRNA4 compared to gRNA1 (n=4 for gRNA1 and n=3 for gRNA4, two-tailed unpaired t-test, p=0.0462). B. Cas9-dependent deletion of miR-124 alters the AMPARs rectification index in CA1 neurons. Schematic representation of the experimental protocol and AAVs used in these experiments (left). Six weeks after co-injection of AAV-Syn-Cre-GFP and either AAV-Flex-Cas9-miR-124^g1^ or AAV-Cas9-miR-124^g4^ in the CA1 of the hippocampus, electrophysiological recordings on GFP^+^ CA1 pyramidal neurons were carried out. The rectification index, a measure of Ca^2+^ permeability of AMPA receptors was lower in the absence of miR-124 (gRNA#4) compared to the control (gRNA#1) (n=18 cells for gRNA#1 and n=15 cells for gRNA#4, 6 mice for each group, two-tailed unpaired t-test, p=0.0294).

To further validate the functional impact of our genome-editing approach, we investigated an additional functional consequence of miR-124 downregulation. Multiple groups have demonstrated that miR-124 is a key miRNA regulating AMPARs composition (Dutta et al., 2013; Gascon et al., 2014; Namkung et al., 2023). Indeed, low levels of miR-124 results in a higher proportion of AMPARs containing the subunit GluA2 encoded by the gene Gria2. Because GluA2-containing receptors are impermeable to Ca^2+^, they exhibit specific channel features that can be assessed by patch-clamp. We therefore explored whether inducing miR-124 deletion in hippocampal neurons *in vivo* with our tools could lead to changes in Ca^2+^ permeability of AMPARs. For that, we constructed new AAVs that enabled Cre recombinase-dependent expression of Cas9 (AAV-Flex-Cas9-miR-124^g4^ and AAV-Flex-Cas9-miR-124^g1^) (Fig 3B). We co-injected these AAVs with an AAV expressing Cre recombinase fused to GFP (AAV-Syn-Cre-GFP) in the CA1 region of the hippocampus. From these *in vivo* injected mice, we then prepared acutely dissected hippocampal slices and performed *ex vivo* whole-cell patch clamp recordings of GFP^+^ CA1 pyramidal neurons (Fig 3B). While stimulating Schaffer collaterals, we recorded AMPARs excitatory post-synaptic currents (EPSCs) at two different membrane potentials (−60 and +60 mV) to measure the rectification index. This electrophysiological measure allows identification of GluA2-containing AMPARs; owing to lack of Ca^2+^-permeability, currents from these receptors show less rectification. As shown in Fig 3B, compared to control AAV (expressing gRNA1), neurons expressing gRNA4 showed a lower rectification index arguing for a higher proportion of GluA2-containing AMPARs. These observations provide an additional confirmation that our tool can disrupt miR-124 expression *in vivo* and lead to functionally-relevant alterations of downstream targets such as Gria2.

## Discussion

In this work, we provide experimental evidence that AAV-mediated delivery of Cas9 can be used to inactivate miR-124 in the mouse brain. We demonstrated that mutations generated at the DNA level resulted in the selective decrease of the mature miRNA levels without affecting the expression of other miRNAs. Using mRNA sequencing in FANS sorted nuclei, we validated the subsequent upregulation of a number of miR-124 target transcripts *in vivo*. More importantly, we confirmed that the reduction of miR-124 levels lead to biological relevant phenotypes. First, we showed that migration of newly generated neurons for the olfactory bulb is perturbed upon miR-124 inactivation. Finally, we also confirmed that manipulating miR-124 with Cas9 resulted in the Gria2-related synaptic modifications providing therefore compelling evidence for the validity of our approach.

Loss of function experiments represent the gold-standard for investigating the function of molecular factors. Although numerous groups have successfully applied Cas9 to target mRNA *in vitro*, this has not been the case *in vivo*. Regarding miR-124, deletion of either mouse miR-124 locus 1 or 3 resulted in a neurodegenerative-like phenotype suggesting maintaining optimal miR-124 levels is necessary for long-term neuronal survival (Liu et al., 2022; Sanuki et al., 2011). Nonetheless, due to compensatory mechanisms, the precise functions of miR-124 could not be studied using these straight knockout mouse lines. Alternative strategies based on microRNAs antagonists (antagomirs, tough decoys or sponges) have been widely used in the past. Nonetheless, this is particularly challenging for highly expressed microRNAs such as miR-124 whose inhibition might require high antagonist levels leading to loss of specificity and/or toxicity. Our work indicated that, using Cas9 for inactivation of endogenous miRNAs, would allow to overcome the above-mentioned technical hurdles. We demonstrated that the net effect of random indels induced by double-strand DNA breaks in the three miR-124 loci is the permanent downregulation of the mature microRNA. In addition, as we showed in Fig 3B, Cas9 could be easily delivered to specific neuronal circuits via loxP-flanked vectors and either injection in transgenic Cre mouse lines or together with a cell-specific Cre AAV.

We provided solid evidence that miR-124 inactivation using our Cas9-AAVs is biologically relevant *in vivo*. We have focused on postnatal neurogenesis in SVZ-RMS-OB. This system is endowed with several advantages to experimentally test the validity of our approach *in vivo*: i) it has been extensively studied; ii) AAVs can be easily delivered intraventricularly to infect progenitor cells in the SVZ; iii) NSCs divide slowly to enable the genetic deletion of the miRNA before diluting the viral vectors; iii) miR-124 function seems to be dispensable for NSCs (Cao et al., 2007; Åkerblom et al., 2012); iv) neuroblasts show high levels of miR-124 and this miRNA has been involved in their migration into a spatially different brain region, the OB (Cheng et al., 2009). Our results provide further support to the notion that miR-124 is essential for the migration of multiple cell types as it has been suggested by previous *in vitro* and *in vivo* work (Cheng et al., 2009; Fang et al., 2014; Li et al., 2013; Volvert et al., 2014).

Finally, we have also confirmed extensive work indicating the importance of miR-124 for regulation of AMPAR composition (Dutta et al., 2013; Gascon et al., 2014; Ho et al., 2014; Hou et al., 2015; Namkung et al., 2023). The balance between Ca^2+^ permeable and Ca^2+^ impermeable AMPARs has been long implicated in multiple brain functions, especially in the cortex (Jonas et al., 1994) and amygdala (Mahanty and Sah, 1998; Clem and Huganir, 2010). Recent data also implicate these receptors in psychiatric diseases (for review see (Jimenez-Gomez et al., 2024; Xu et al., 2024)). A remarkable example of the physiological relevance of Ca2+ permeability of AMPARs has been recently shown in the visual cortex (Hong et al., 2024). Thus, it has been long known that pyramidal neurons express calcium-impermeable AMPARs and are very selective to precise stimuli. In contrast, parvalbumin interneurons express AMPARs lacking the Gria2 subunit (calcium-permeable AMPARs) and are characterized by a low stimuli selectivity. Hong et al demonstrated that experimental manipulation of Gria2 expression in these different cell types and therefore switching from one type of receptors to the other affected their orientation tuning. Since miR-124 tightly controls Gria2 expression and its expression is activity dependent (Krol et al., 2010; Sambandan et al., 2017), we suggest that miR-124 might orchestrate functional refinement in complex neuronal networks by finely regulating AMPA receptors in different cell types according to their actual inputs.

Although our findings are promising, there are some caveats that would need to be addressed by future work. Similar to knockout mice, compensatory mechanisms might be also activated in response to Cas9-induced deletion. Thus, for example, primary transcript abundance (pri-miR-124) is increased in Cas9^+^ cells likely reflecting enhanced transcription and/or decreased processing (Fig 1C). Although the functional impact of such increase remains unknown, results from plant miRNAs (Lauressergues et al., 2015) and long non-coding RNAs (Zhou et al., 2022) suggest that functional short regulatory peptides might be encoded from these transcripts. A second issue of our approach concerns the insertion of random mutations at highly expressed genomic loci. This intrinsic feature of DNA repair machinery ensures miR- 124 loss while limiting the likelihood of creating novel miRNA species. Nonetheless, it is impossible to completely rule out that none of the mutations is functional. Given the abundance of miR-124 in the brain, the relevance of such event should not be underestimated. Finally, our FANS sorting approach enables the isolation and specific analysis of infected cells in which genome editing is possibly occurring. However, there is no cell specificity in our AAV vectors as CMV promoter has been reported to be active in multiple neuronal types in the cortex (Watakabe et al., 2015). Consequently, our Cas9^+^ nuclei fraction likely encompasses a heterogeneous neuronal population. Since transcripts under miR-124 regulation might widely vary among neuronal subsets (Parkins and Gross, 2024), our mRNA sequencing findings might have uncovered only those shared in enough cells, which might represent only a small fraction of mRNAs actually deregulated. Future experiments using single nucleus RNA sequencing are crucial to address this issue and to identify cell-specific miR-124 targets.

In summary, our findings support the remarkable efficiency (strong downregulation of miR-124), specificity (no off-targets could be detected and no other miRNAs are affected) and physiological relevance (downstream biological processes altered) of our new Cas9-tools. Since we showed that Cas9-tools could be packed in AAVs vectors and coupled to Cre-loxP system, our work therefore opens new avenues for interrogation of miRNA functions in specific neuronal networks *in vivo*.

## Methods

### gRNA designs and plasmids

Guide-RNAs targeting miR-124 were manually designed using the mature sequence of the microRNA as well as 25 bp upstream/downstream to identify potential PAM sites for saCas9. The sequences of the 4 tested guides can be found in table 2. For cloning of gRNAs into a plasmid containing saCas9 (Addgene # 61591), we followed the recommended protocol from Zhang’s lab (available on Addgene). For transfection in cell lines expressing a miR-124 reporter, we further modified this plasmid by adding a PGK-CFP cassette at the NotI site (from a modfied pSuper neo+GFP, Oligoengine). For CA1 hippocampal injections, gRNA1 and gRNA4 were cloned into an analogous vector but in which saCas9 is in a Flex configuration (pAAV-FLEX-SaCas9-U6-sgRNA, Addgene #124844). A pAAV-Syn-Cre-GFP plasmid was also used for viral production (Addgene #105540).

**Table 2.**
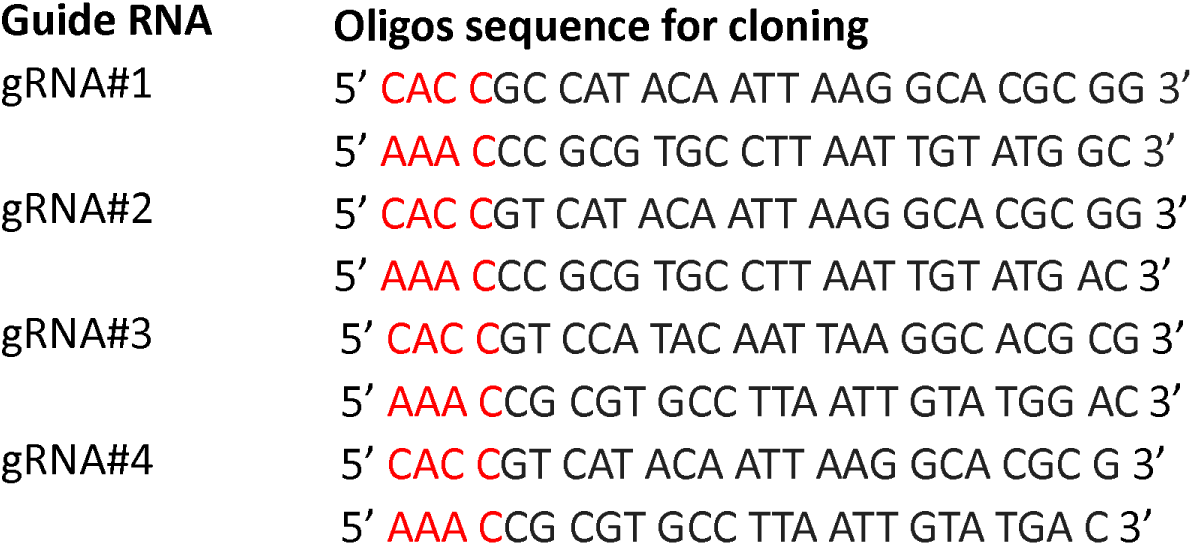
List of gRNAs tested in this work as well as the oligos used for cloning.

### Cell culture and transfection

HEK293T cells were obtained from American Type Culture Collection (ATCC) and cultured under standard conditions (DMEM-10% Fetal Calf serum). For transfection, we used Lipofectamine 3000 (Thermo) as transfection reagent following manufacturer’s recommendations.

### Adeno-associated vector production

Viral productions were carried out using standard protocols with slight modifications (Su et al., 2020). Briefly, HEK293T cells were transfected using the calcium phosphate method (pHelper, pRC2, pAAV; ratio 2:1:1). After 60-72h, cells were harvested and AAV particles purified and concentrated using Takara dedicated kit (Takara, France). Viral productions were tittered using quantitative PCR (Takara, France). AAV9 serotypes of a titer around 10^12^ viral genomes/ml were used in this work. The different vectors are listed in Table 3.

**Table 3.**
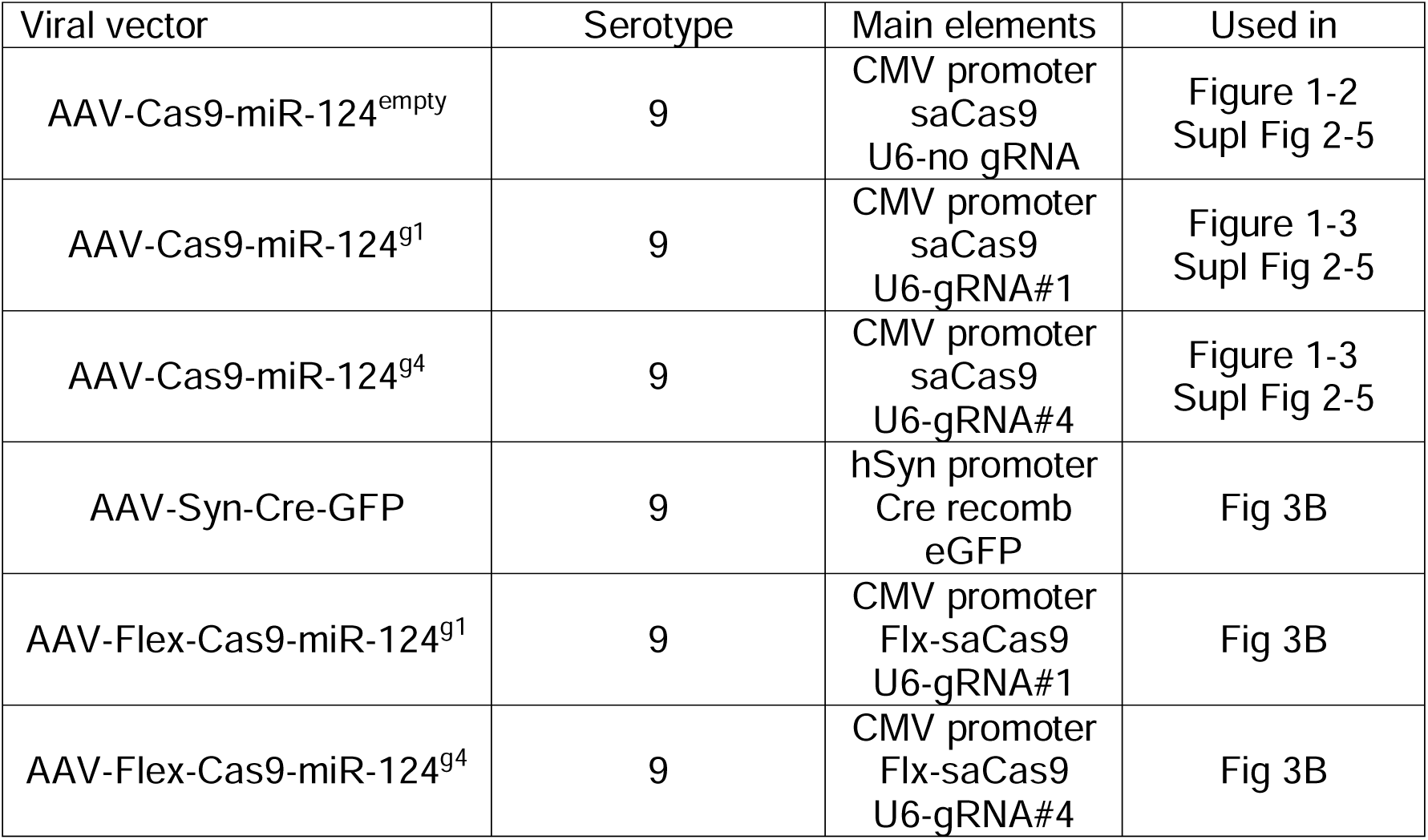
List of viral vectors used in this work.

### Animals

All mice were purchased to Charles Rivers (France). C57Bl6J (9-10 weeks) mice were kept in the local animal facility, under controlled temperature and a 12 h/12 h light-dark cycle (lights on at 8 AM to 8 PM). Animals had access to food and water *ad libitum*. All procedures have been approved by the local ethics committee and are in agreement with European regulations (Directive 2010/63/EU). A special effort was made in handling animals to reduce the number of animals as well as to minimize stress or anxiety. A total of 34 mice was used in this work.

### Surgical procedures

Mice were deeply anesthetized with ketamine (100 mg kg^−1^) and xylazine (10 mg kg^−1^) or by breathing isofluorane (2-4%) through a mask. Then, they are a placed in a stereotaxic frame. The coordinates according to the mouse brain atlas of Paxinos & Franklin were as follows: for the motor cortex (M2) AP, 1,4 mm; ML, ± 1 mm; for hippocampal CA1 AP, −2,2 mm; ML, ± 1.5 mm relative to the bregma. For DV, we used −0,4 mm for cortical injections relative to the brain surface and −1,5 mm for the CA1 relative to the skull. We injected high-titer vector (0.2-1.5 × 10^13^; 1 μl). For M2 injections, we used either AAV-Cas9-miR-124^empty^, AAV-Cas9-miR-124^g1^ or AAV- Cas9-miR-124^g4^ (two injections/hemisphere). For CA1 injections, animals were injected with a mixture of AAV-Syn-Cre-GFP and either AAV-Flex-Cas9-miR-124^g1^ or AAV-Flex-Cas9-miR-124^g4^. Injections were carried out using a microinjector (World Precision Instruments) at 333 nL/min (motor cortex) or a micromanipulator (Narishige).

The intracerebroventricular injection protocol was adapted from previous work (Leprince et al., 2023). Briefly, mouse pups were anesthetized by hypothermia and were mouth-injected into the lateral ventricles the viral solution using a glass pipette pulled from borosilicate capillary tubes. To reach the ventral horn of the lateral ventricle, we drew an imaginary line between lambda and the eye and injected at about 1/3 of this line and at a depth of 0.3 mm. We filled the right ventricle with 1,5-2 μL of a mixture of AAV-Cas9-miR-124^g1^ or AAV-Cas9-miR-124^g4^. Ten days later, animals received 4 intraperitoneal injections of the thymidine analog, BrdU, (50 mg/kg, Sigma, diluted in saline) every 2 hours to label actively proliferating cells.

### Cytometry and FANS analysis

For nuclei sorting, all steps were performed at 4 °C or on ice. Tissues were homogenized in nuclei isolation buffer (0.32 M Sucrose, 10 mM HEPES pH 8.0, 5 mM CaCl_2_, 3 mM Mg (CH_3_COO)_2_, 0.1 mM EDTA, 1 mM DTT, 0.1% Triton X-100) with a 2 ml Dounce homogenizer by 30 strokes with each pestle and filtered through a 70 µm strainer. After centrifugation, nuclei pellets were resuspended in 1 ml PBS-RI (PBS, 50 U/mL Rnase-OUT Recombinant Ribonuclease Inhibitor (Invitrogen), 1 mM DTT) and fixed by the addition of 3 ml PBS 1.33% paraformaldehyde (Electron Microscopy Sciences) for 20 minutes. Fixed nuclei were spun down, washed with 1 mL PBS-RI 0.1% triton-X-100, pelleted again and resuspended at 10^6^ nuclei per mL in stain/wash buffer (PBS-RI, 0.5% BSA, 0.1% Triton-X-100) containing 20 µL/mL anti-HA-tag-alexa-647 antibody (R&D Biosystems #IC6875R) and 1 µg/mL Hoechst 33342 (Molecular Probes). After 30 minutes incubation, nuclei were washed with 2 mL stain/wash buffer and spun down. Finally, stained nuclei were resuspended in 1 ml PBS-RI 0.5% BSA and filtered again through a 40 µm strainer. Nuclei suspensions were maintained on ice protected from light until sorting.

Sorting of nuclei was achieved with a Cytoflex SRT cell sorter (Beckman Coulter). After positive selection of intact Hoechst-positive nuclei and doublets exclusion, all HA-positive and 2000 HA-negative nuclei were separately isolated. Sorted nuclei were collected in 2 ml microtubes pre-coated with BSA and finally were conserved at −80°C until RNA extraction.

For cell cytometry *in vitro*, transfected cells were trypsinized, rinsed in PBS and resuspended in the same buffer as for sorting. Fluorescence was detected using a Cytoflex device.

### Analysis of miR-124 genomic edition

We first used DNA sequencing for analyzing mutations at the DNA level. Sorted nuclei from injected M2 were recovered into direct lysis buffer (TaqMan™ Fast Advanced Cells-to-CT™ Kit, ThermoFisher Scientific). From the possible off-targets provided by Cas-OFFinder (Bae et al., 2014), we focused on the sequences showing the highest similarity to those of the on-targets (16 nt out of 21): 7 sequences for the gRNA4 guide, 3 sequences for the gRNA1 guide and 1 sequence common to both. We pooled the samples from the same experimental group (empty/HA^-^, g1/HA-, g4/HA^-^, empty/HA^+^, g1/HA^+^, g4/HA^+^) and amplified the 16 loci from 1 μL of pool (primers in Table 4). We first amplified the sequences and purified the amplicons using AMPure XP Beads (Beckman Coulter), with a buffer/sample ratio of 0.4 and then 1.8. We pooled amplicons for each experimental group of final 6 pmol. Then we performed A-tailing with the Klenow enzyme and cleaun-up with AMPure XP Beads, ratio 1.8. For library generation, we used QIAseq 1-Step Amplicon Library Kit (Qiagen, ref. 180419) starting with 80 ng of sample and following manufacturer guidelines. Libraries were purified with AMPure XP Beads, first with a buffer/sample ratio of 0.4 and then 1.8. We pre-amplified the libraries with HiFi PCR Master Mix as recommended and purified them with AMPure XP Beads, ratio 1. The libraries were sequenced on an MiniSeq device (Illumina).

**Table 4.**
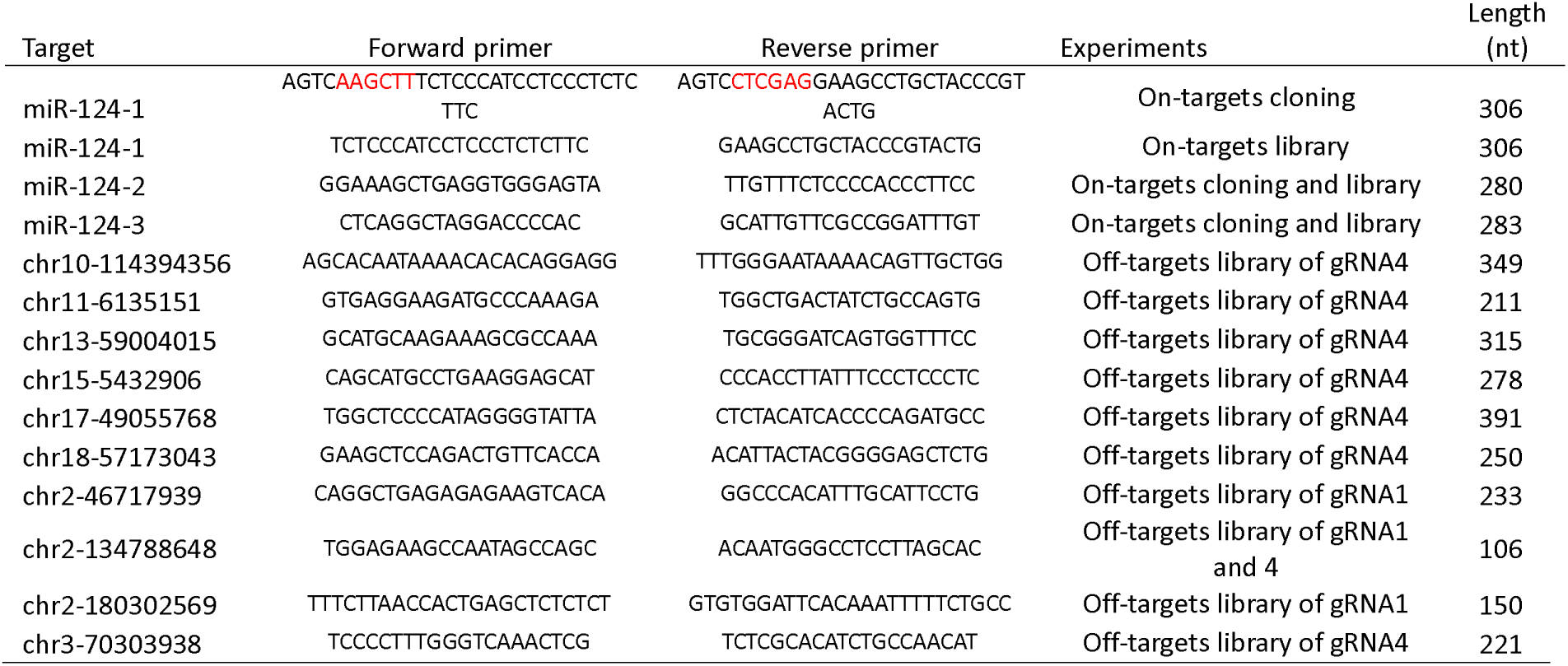
List of primer sequences used for DNA sequencing.

The reads were ordered based on the type of mutation (substitution, insertion, deletion). As substitutions were observed along all sequences in a random manner, substitutions and indels (insertions + deletions) were analyzed separately. For each position, we calculated the percentage of times that a position appeared mutated with respect to a reference genome (*Mus musculus*). Readings with less than 1% cases were not taken into account.

To validate the effect of the gRNA, we also performed Sanger sequencing of some experimental groups after cloning the amplicons of loci 1, 2, and 3. For locus 1 we used the pCRII vector previously digested with HindIII and XhoI (ThermoFisher Scientific). We inserted these sites in the primers of locus 1 (Table 4). For loci 2 and 3 we used the pCR4-TOPO vector (TOPO-TA cloning kit, ThermoFisher Scientific) (primers Table 4). Bacterial transformation and selection, plasmid purification using MiniPreps (Machery Nagel) and Sanger sequencing (Eurofins genomics) were performed using standard protocols as described before (Gascon et al., 2014).

### Total RNA extraction

We first removed the cross-linker incubating 30 min at 70 °C in TE buffer as previously described (Evers et al., 2011) Total RNA was then purified using Qiazol Lysis Reagent (Qiagen #79306). Briefly, 800 µL of Qiazol were added to the sorted nuclei with 1 µL UltraPure Glycogen (20µg/µL, ThermoFisher Scientific #10814010) and 160 µL chloroform. After centrifugation, the upper layer was retrieved and RNA was precipitated, washed and finally resuspended with 20 µL TE buffer. RNA quality and quantity were determined on a BioAnalyzer with RNA 6000 Pico reagents and chips (Agilent #5067-1513).

### miRNA and pri-miRNA quantification by qPCR

Mature miRNAs were reverse-transcribed with the TaqMan Advanced miRNA cDNA Synthesis kit (ThermoFisher Scientific, #A25576). 3,7 µL of purified total RNA were used as template for poly-adenylation followed by 3’ adapter ligation, reverse transcription and 14 cycles of pre-amplification.

For pri-miRNA quantification, 5 µL of total RNAs were reverse transcribed with SuperScript IV reverse transcriptase and random hexamers (ThermoFisher Scientific), then cDNAs were pre-amplified with a 0.05X pool of TaqMan assays for 14 cycles.

Pre-amplified cDNAs were diluted 1:10 with 0.1X TE buffer. 2.5 µL of this dilution was used as template in quantitative PCR reactions with 1 µL of TaqMan assay (ThermoFisher Scientific), 5 µL of TaqMan Fast Advanced Master Mix (ThermoFisher Scientific) and 2 µL water. qPCR reactions were run on a QuantStudio7 Flex system (ThermoFisher Scientific).

### RNA sequencing

40 to 2000 pg of purified total RNA were used as starting material for the preparation of libraries using the QIAseq FastSelect RNA library kit (Qiagen, #334235). First, possible DNA contamination was removed by DNAseI treatment (Amplification grade DNaseI, ThermoFisher Scientific, # 18068015). Then, following the Qiagen kit protocol, rRNA was depleted, then cDNA was reverse-transcribed using both oligodT and random hexamers and a barcode was added to each sample. Finally, cDNA was amplified (27 cycles) and indexed with QIAseq UX 96 Index Kit IL UDI-B (Qiagen # 331825). The libraires were equimolarly pooled and sequenced on a NextSeq instrument (Illumina) for 74 bp paired-end and dual 10 bp index reads.

### In situ hybridization and immunofluorescence

Adult mice injected with Cas9 AAVs in the cortex or young animals having received BrdU were deeply anesthetized with ketamine (100 mg kg^−1^) and xylazine (10 mg kg^−1^) and perfused intracardially with ice-cold PBS followed by 4% paraformaldehyde. Brains were extracted, postfixed in the same fixative overnight at 4 °C, transferred to a 30% sucrose solution for cryoprotection, frozen, and stored at −80 °C. We sectioned brain samples with a standard cryostat (Leica 3050). 20 μm sections were directly transferred to a microscope slide for in situ whereas 40 μm floating sections were used for immunostaining

Protocols used in this work are standard and have been described elsewhere (Gascon et al., 2007; Gascon et al., 2014). For detection of miR-124, we purchased 5’ and 3’ digoxigenin-labeled LNA probes from Exiqon. Briefly, after proteinase K treatment (10 min at 37 C), sections were extensively rinsed and preincubated for 3h at room temperature with the hybridization buffer (50% formamide, salmon sperm DNA, tRNAs). Probe was hybridized overnight at 55 °C, and the slides were incubated with horseradish peroxidase-conjugated antidigoxigenin antibody (Roche; 1:500) and steptavidin linked to the peroxidase (Roche). Cy3 TSA Plus kit (Perkin-Elmer) were used for final detection.

For immunofluorescence, we diluted primary antibodies in PBS containing 10% donkey serum (Sigma), 3% bovine albumin (Sigma), and 0.3% Triton X-100 and incubated overnight at 4 °C. Corresponding donkey anti-rabbit or anti-mouse Alexa 488 or 555 conjugated secondary antibodies (Invitrogen) were used for secondary detection (2h at room temperature, dilution 1/1000 in PBS). Before being mounted, sections were counterstained with Hoescht (ThermoFisher Scientific) (5 min at room temperature, dilution 1/5000). For BrdU detection, slices were incubated with 2 M HCl and rinsed in PBS before the reaction with the primary antibody. Primary antibodies used were mouse monoclonal anti-NeuN (Millipore, MAB377) (1:1000 dilution), monoclonal rat anti-BrdU (Abcam, ab6326) (1:500 dilution) and rabbit polyclonal anti-DCX (Abcam, ab18723).

### Electrophysiology in hippocampal slices

For ex vivo electrophysiology recordings, mice were culled by cervical dislocation six weeks after AAV injection. Hippocampi were dissected and incubated for 5 min in ice-cold oxygenated (95 % O_2_/ 5 % CO_2_) cutting solution (in mM): 234 sucrose, 2.5 KCl, 1.25 NaH_2_PO_4_, 10 MgSO_4_, 0.5 CaCl_2_, 26 NaHCO_3_, 11 glucose (pH 7.4). Hippocampal slices (250 μm) were cut on a vibratome (Microm HM600V, Thermo Scientific, France). For recovery, slices were then incubated in standard aCSF (in mM): 119 NaCl, 2.5 KCl, 1.25 NaH_2_PO_4_, 26 NaHCO_3_, 1.3 MgSO_4_, 2.5 CaCl_2_ and 11 D-glucose, oxygenated with 95 % O_2_ and 5 % CO_2_, pH 7.4 for 1 h at 37 ± 1 °C and then stored at RT until used for recordings. Recordings were done in this standard aCSF in a recording chamber on an upright microscope with IR-DIC illumination (SliceScope, Scientifica Ltd, UK) using a Multiclamp 700B amplifier (Molecular Devices, San Jose, CA, USA), under the control of pClamp10 software (Molecular Devices, San Jose, CA, USA). Whole-cell patch-clamp recordings of CA1 pyramidal neurons were performed at 31 ± 1°C. The Schaffer collateral pathway was stimulated at 0.1 Hz using electrodes (glass pipettes filled with aCSF) placed in the stratum radiatum. The recording pipette (5-6 MΩ) was filled with a solution containing the following: 117.5 mM Cs-gluconate, 15.5 mM CsCl, 10 mM TEACl, 8 mM NaCl, 10 HEPES, 0.25 mM EGTA, 4 mM MgATP and 0.3 NaGTP (pH 7.3; osmolarity 290-300 mOsm).

After a tight seal (>1GΩ) on the cell body of the selected neuron was obtained, whole-cell patch clamp configuration was established, and cells were left to stabilize for 2-3 min before recording began. Holding current and series resistance were continuously monitored throughout the experiment, and if either of these two parameters varied by more than 20%, the cell was discarded. Excitatory post-synaptic currents (EPSCs) were pharmacologically isolated by adding 50 μM picrotoxin (Sigma-Aldrich, dissolved in DMSO) to block GABAergic transmission and 50 μM APV (Tocris, dissolved in water) to block NMDA transmission.

### Image acquisition

All photographs were acquired on an Axiovert (Zeiss) equipped with a digital camera (Hamamatsu). To ensure no differences in signal intensity, digital images of olfactory bulbs (5-10 for each animal) were acquired with fixed settings.

### Data analysis

For *in vitro* experiments, qPCR was analyzed using miR-92 as reference. For quantification of reporter intensity, the ratio between median RFP signal and CFP (internal reference) was calculated

For RNA-sequencing, data were demultiplexed with bcl2fastq (Illumina, v2.20.0.422) and read quality was assessed with fastQC (0.11.5). Raw sequencing data in FASTQ format were pre-processed using the GeneGlobe platform (https://geneglobe.qiagen.com/fr/analyze). This step included quality control, adapter trimming, and alignment to the mouse reference genome (Mus musculus GRCm38.101). Then raw count data representing the number of sequencing reads mapped to each gene were exported from GeneGlobe for further differential gene expression (DGE) analysis using the DESeq2 (1.44.0) package in Rstudio (4.4.1). Among the genes that had at least 1 read (Hapos: 17,570 genes ; Haneg : 18,117 genes), genes with more than 160 counts were used (Hapos : 5,483 genes ; Haneg : 4,822 genes) to run the DESeq analysis for comparison of the experimental conditions (g4 vs Empty). As the vast majority of genes had a similar expression in the 2 groups, only the 660 genes with the highest FC values (HApos |FC| > 1.5 ; Haneg |FC| > 1.33) were conserved and the adjusted p-value was obtained with the FDR method.

For DNA sequencing, the raw output of illumina (https://www.illumina.com/) binary base call files (BCL) were treated and demultiplexed with bcl2fastq (https://emea.support.illumina.com/sequencing/sequencing_software/bcl2fastq-conversion-software.html) to produce the fastq files for each of the CAS9 experimental condition. Then, each of these fastq files were further trimmed using the cutadapt (https://github.com/marcelm/cutadapt?tab=readme-ov-file) to produce the corresponding preMiR sequences of fastq files. Each sequence from each of these demultiplexed preMir sequence files were aligned (pairwise alignment) to their corresponding reference sequence using blastn (Camacho et al., 2009). The number of mutations in the alignments (https://evolution.berkeley.edu/dna-and-mutations/types-of-mutations/) were counted only if the alignment length was, at least, equal to the length of the reference sequence. Each case of 3 types of mutations namely insetion, deletion and substitution are taken into account when quantifying mutations. Thus, a count table of mutations for each demultiplexed sequence files were created and exported to tabular file for further analysis.

For quantification of BrdU nuclei in olfactory bulb slices, at least, 10 sections/hemisphere from 7 different animals (4 injected with AAV-Cas9-miR-124^g1^ and 3 injected with AAV-Cas9-miR-124^g4^) were automatically quantified using ImageJ and the cell counter tool. We then calculated the ratio of BrdU^+^ nuclei between the right (injected) and left (non-injected) sides that was used for comparisons.

For electrophysiological recordings, data was acquired using Clampfit 10 software (Molecular Devices, San Jose, CA, USA). For rectification index evoked EPSCs recording, neurons were voltage clamped first at −60 mV, recorded every 10 s for a minimum of 15 sweeps and then clamped at + 60mV for a minimum of 15 sweeps. Amplitude mean per voltage was calculated and the rectification index was calculated as the mean EPSC −60mV/mean EPSC +60mV. The experimenter was blind as to the condition recorded and analyzed.

### Statistical analysis and experimental design

Detailed statistical procedures for each experiment are provided in each figure legend. For *in vitro* experiments, all the conditions were run simultaneously and replicated at least 3 times. Data were analyzed with Prism and statistical differences using 2 or 3-way ANOVA and post-hoc tests.

For *in vivo* experiments, pools of animals (DNA sequencing) or individual animals injected on one hemisphere with AAV-Cas9-miR-124^empty^ or AAV-Cas9-miR-124^g4^ (qPCR and RNA sequencing) were used. Data were statistically analyzed using FDR (DEseq2 R package) for RNA sequencing or 2-way ANOVA and Sidak post-hoc test (Prism) for qPCR.

For functional experiments (BrdU and electrophysiology) data analysis were performed on Prism using unpaired t-test.

## Data Availability Statement

All data supporting the conclusions are available via the corresponding author upon reasonable request. Sequencing data are deposited on NCBI’s Gene Expression Omnibus (Edgar et al., 2002) and are accessible through GEO. For RNA-sequencing, GSE291994 (https://www.ncbi.nlm.nih.gov/geo/query/acc.cgi?acc=GSE291994) and GSE291957 (https://www.ncbi.nlm.nih.gov/geo/query/acc.cgi?acc=GSE291957) for DNA- sequencing.

## Author Contributions

DRM, NP, FJ, CR, CL, AB, MV and SM performed all the experiments and helped out in the writing of the first draft of the manuscript. DB, DRM, NP: bioinformatics analysis. HM: electrophysiological experiment design and analysis. MC: design of newborn experiments and writing; EG: writing, review and editing, supervision, project administration and funding acquisition. All authors contributed to the article and approved the submitted version.

## Funding

This work was supported by the French National Agency (ANR) to EG (ANR-22- CE17-0034), Fondation de France (00100077), Fondation pour la recherche sur Alzheimer (AAP2022) and Fondation France Alzheimer (2021-#6239) to EG.

## Conflict of Interest

The authors report no biomedical financial interests or potential conflicts of interest.

## Acknowledgements

The authors would like to thank NeuroVir viral core for the viral productions and Stephane Robert from AMUTICYT facility for technical support with cytometry and FACS.

## Notes

### Competing Interest Statement

The authors have declared no competing interest.

