## Supplementary material for "Combination of Cas9 and adeno-associated vectors (AAVs) enables efficient *in vivo* knockdown of precise miRNAs in the rodent brain": Supplemetary figures 1-5

#### Roura-Martinez et al Suppl Figure 1

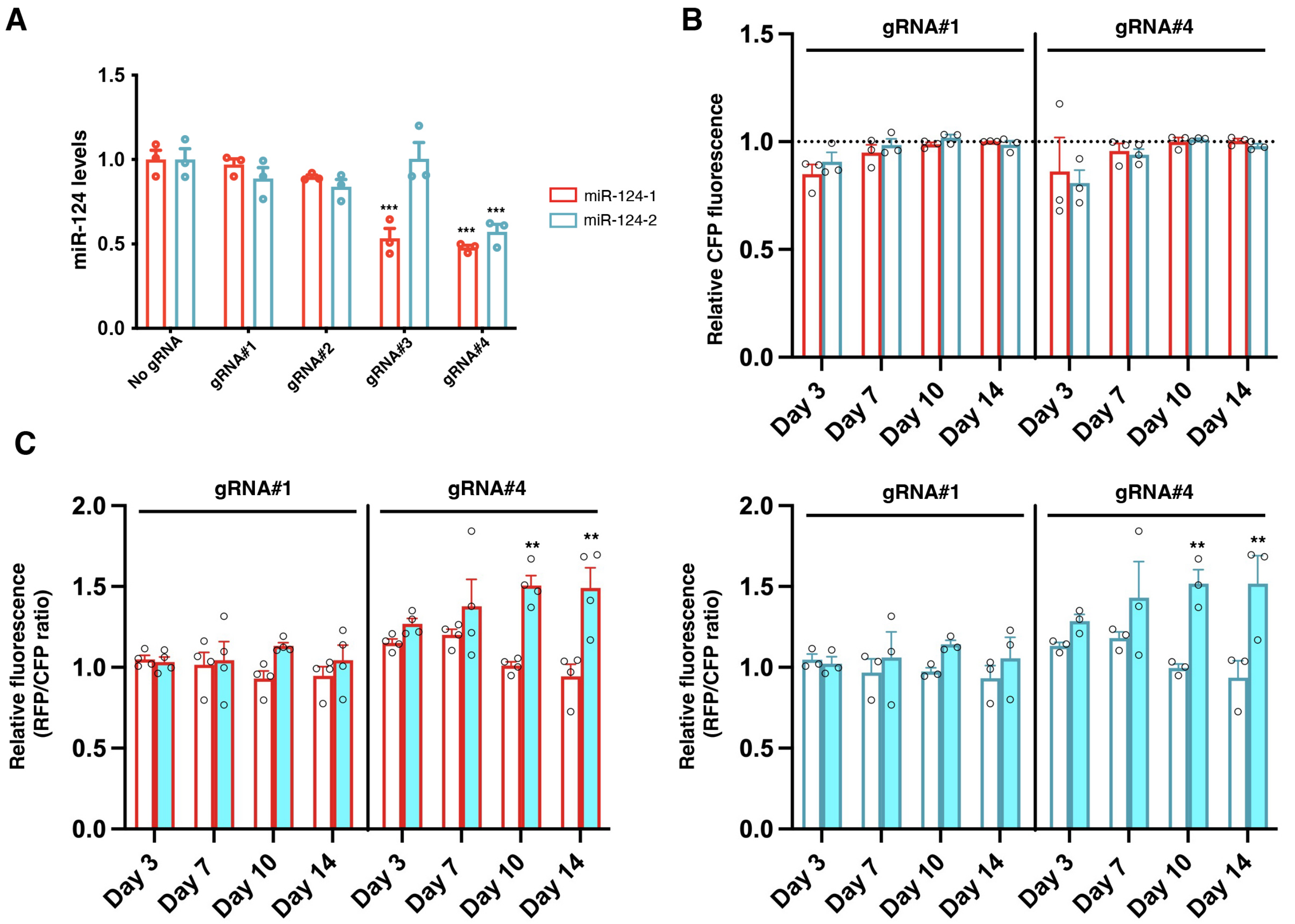


Supplementary Figure 1. Cas9-mediated inactivation of miR-124 in vitro

A. HEK stably expressing either mouse miR-124-1 or miR-124-2 were transfected with a plasmid driving the expression of saCas9 and one of the gRNAs to be tested. miR-124 levels were quantified 10 days later using qPCR (miR-92 was used as endogenous control). Graphs represent the results of n=3 independent experiments (2-way ANOVA, $F(4,20)=22.31$, $p<0.0001$). Multiple comparison for miR-124-1 : gRNA1 vs cont p=0.9844, gRNA2 vs cont p=0.5, gRNA3 vs cont p<0.0001, gRNA4 vs cont p<0.0001 ; for miR-124-2 : gRNA1 vs cont p=0.4248, gRNA2 vs cont p=0.1556, gRNA3 vs cont p>0.9999, gRNA4 vs cont p<0.0001(Dunnett's post-hoc test).

B. Cas9/gRNA expression does not interfere with the expression of a control CFP reporter. Cells expressing both miR-124 and a RFP reporter containing binding sequences for miR-124 were transfected with a plasmid driving the expre ssion of saCas9, gRNA1 or 4 and a control CFP reporter (no regulation by miR-124). As shown in the graphs, CFP fluo rescence was not modified at any time point. Values were normalized against cells expressing the same CFP reporter but no Cas9. Graphs represent the results of n=3 independent experiments (2-way ANOVA, $F(1,8)=0.487$, $p=0.505$).

C. Inactivation of miR-124 increase the fluorescence of a miR-124 dependent reporter. In the same experiments as abo ve, the ratio between RFP and control CFP was determined using cytometry at different time points. Compared to gRNA1, cells expressing gRNA4 showed an increase in this ratio starting 10 days post transfection. Graphs represent the results of n=3 independent experiments (3-way ANOVA, $F(1,28)=22.64$ $p<0.0001$ for miR-124-1 and $F(1,28)=21.91$, $p<0.0001$ for miR-124-2). Multiple comparison for miR-124-1 (CFP- vs CFP+): 3 days: gRNA1 p=0.9999 gRNA4 p=0.9666 7 days: gRNA1 p=0.9999 gRNA4 p=0.6395, 10 days: gRNA1 p=0.4225 gRNA4 p=0.0001, 14 days: gRNA1 p=0.9938 gRNA4 p<0.0001; for miR-124-2 (CFP- vs CFP+): 3 days: gRNA1 p=0.9999 gRNA4 p=0.9182, 7 days: gRNA1 p=0.9989 gRNA4 p=0.3613, 10 days: gRNA1 p=0.8610 gRNA4 p=0.0019, 14 days: gRNA1 p=0.9823 gRNA4 p=0.0006 (Tukey's post-hoc test).

#### Roura-Martinez et al Suppl Figure 2

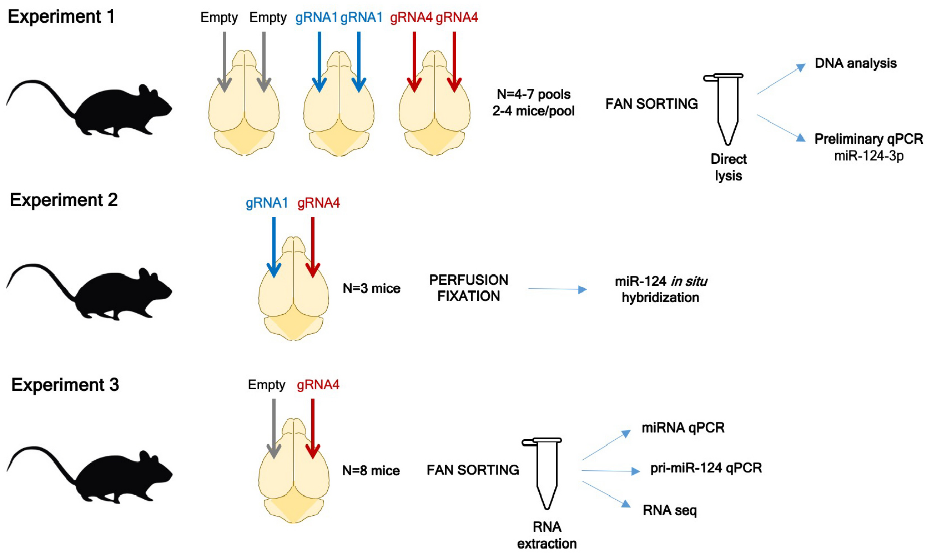


Supplementary Figure 2. Experimental designs of in vivo experiments in the mouse motor cortex.
Different combinations of AAVs depicted in Fig 1A were injected into the motor cortex of adult mice. To examine miR-124 inactivation, after four to six weeks, mice were sacificed and used for the indicated downstream applications.

### Roura-Martinez et al Suppl Figure 3

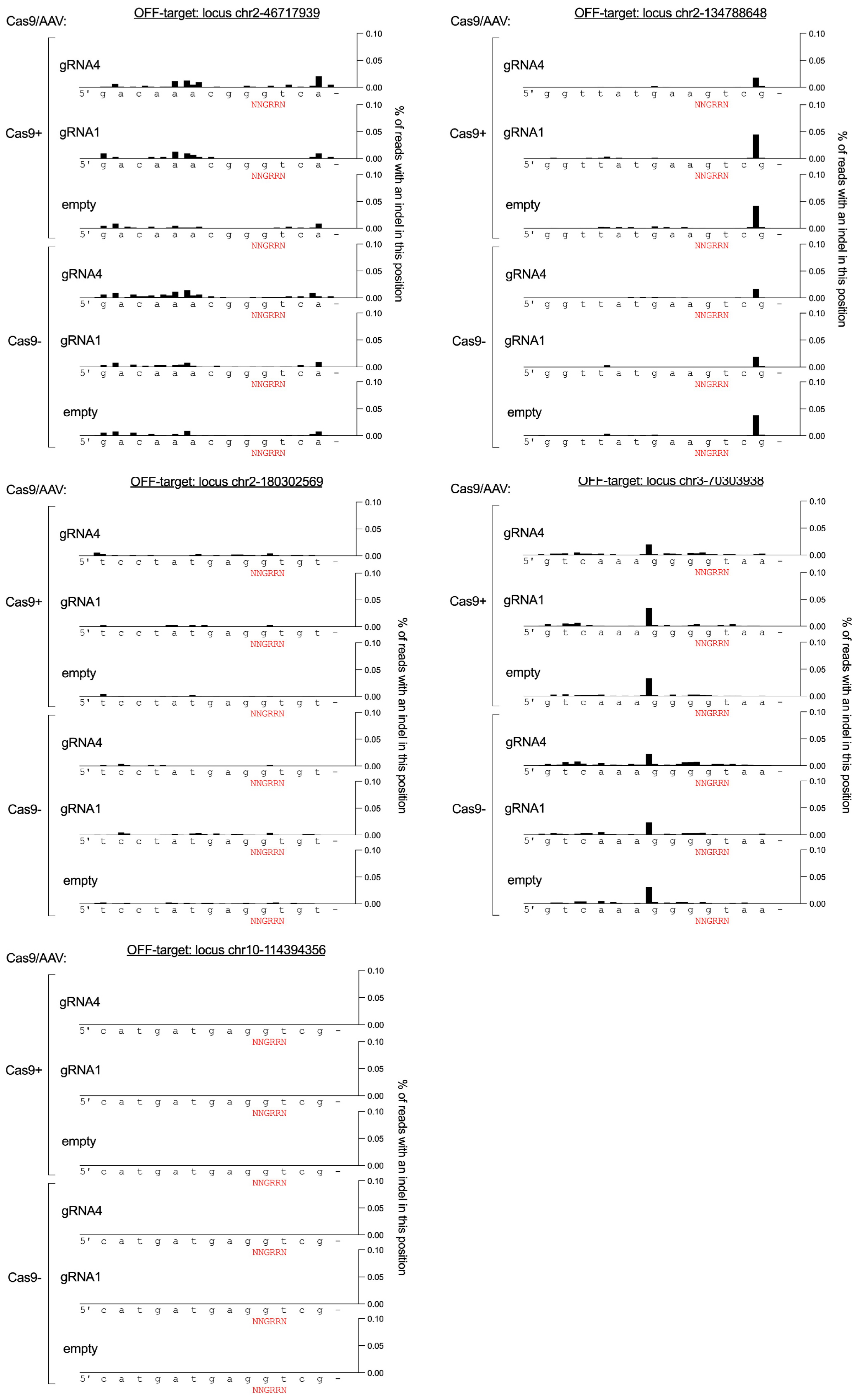

### Roura-Martinez et al Suppl Figure 3

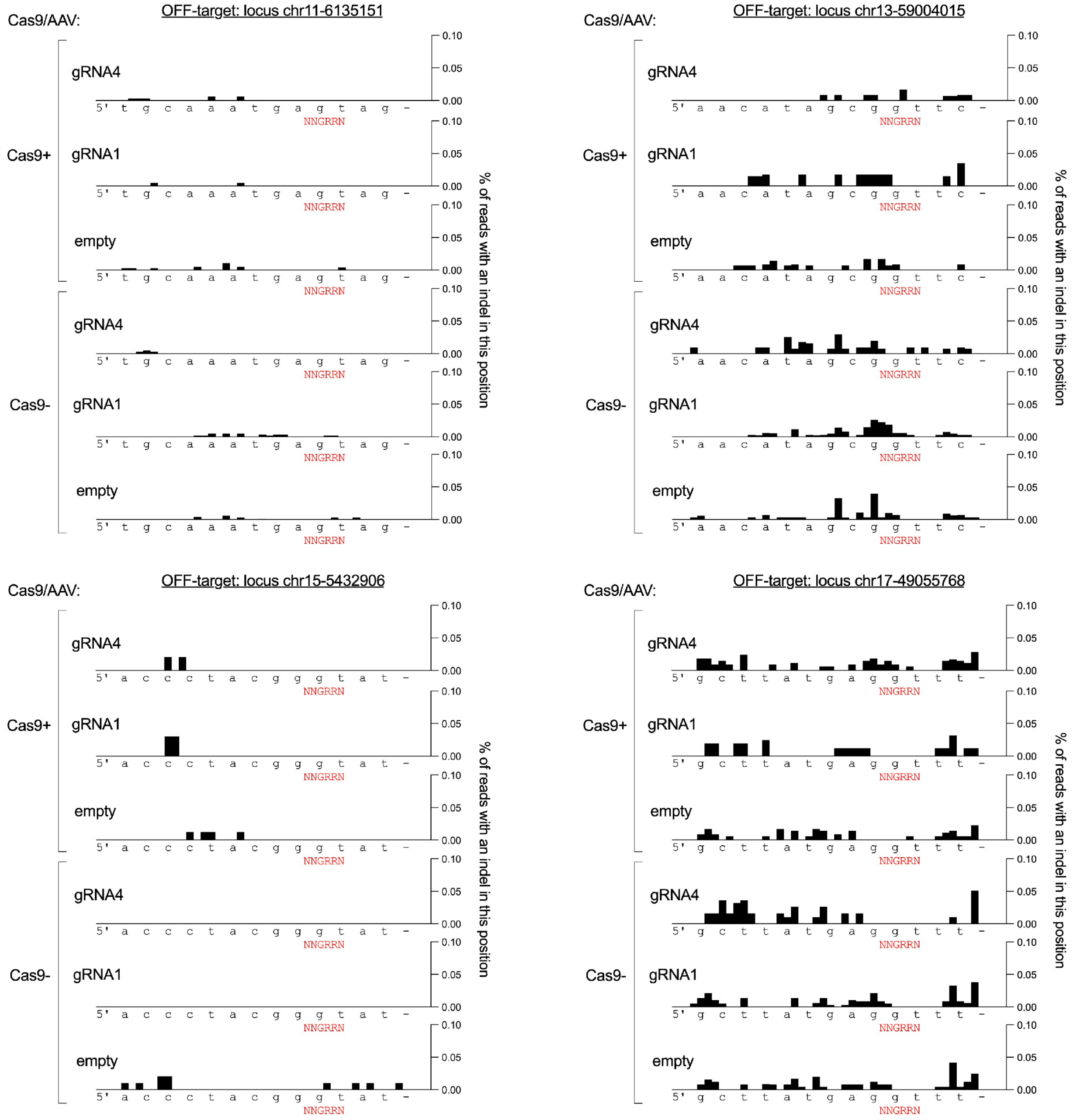


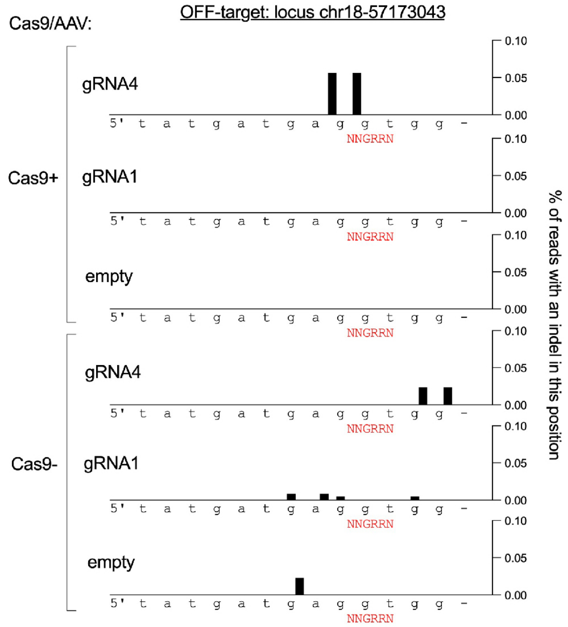


Supplementary Figure 3. Evaluation of Cas9 off targets in vivo. Using DNA sequencing, we analyzed potential mutation in the 10 top off target loci for gRNA4. As shown in the graphs, no signi ficant differences in the mutation rate were observed among the conditions (AAV-miR-124-Cas9$^{empty}$, AAV-miR-124-Cas9$^{g1}$ and AAV-miR-124-Cas9$^{g4}$) indicating that the occasional mutations found might reflect technical noise from DNA amplification.

**Roura-Martinez et al Suppl Figure 4**

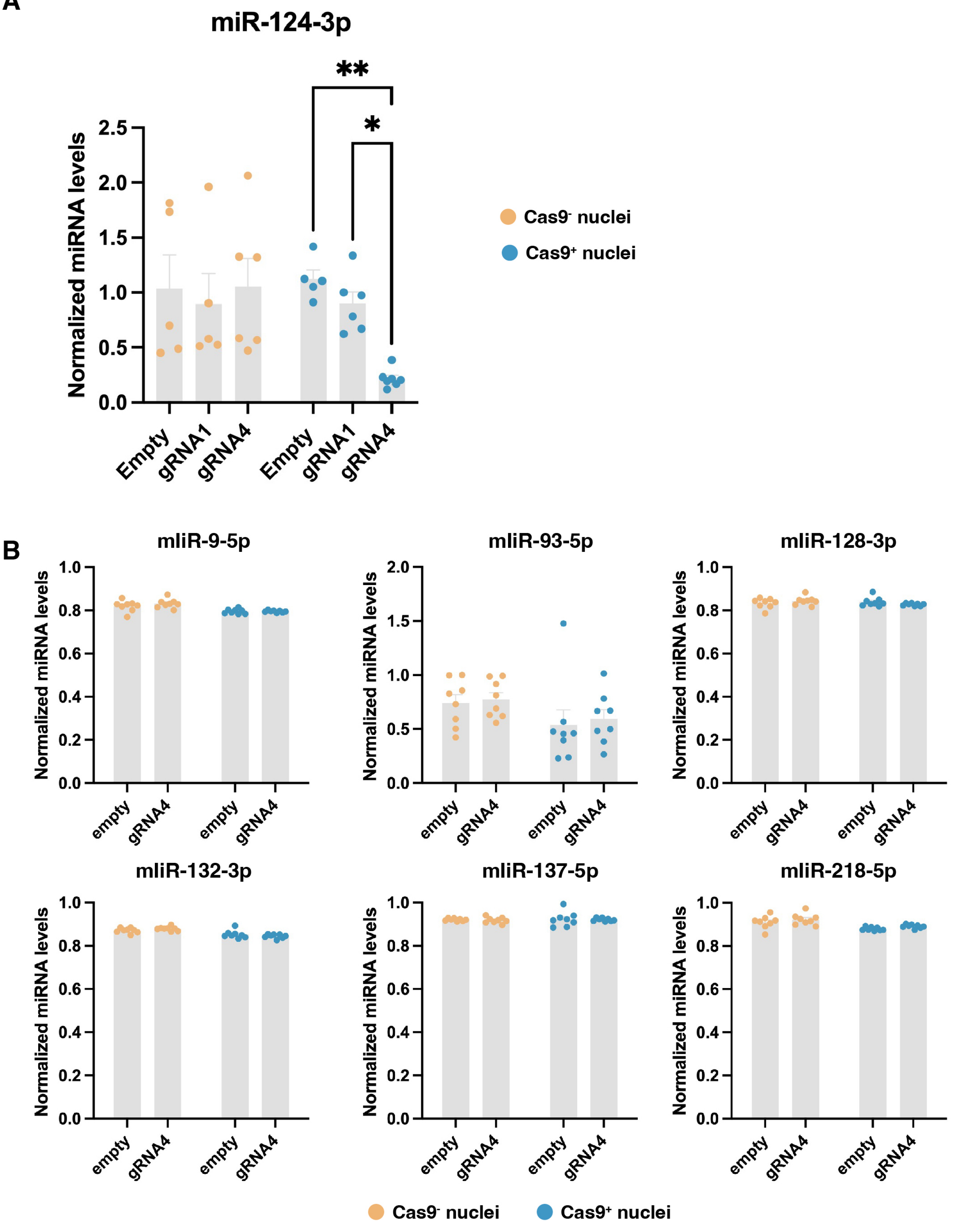


Supplementary Figure 4. Additional control for Cas9 experiments in vivo.
A. miR-124-3p expression was quantified in pools of animals injected with AAV-miR-124-Cas9$^{g4}$ or any of the control vectors (AAV-miR-124-Cas9$^{empty}$ or AAV-miR-124-Cas9$^{g1}$). Graphs represent the results of n=5-7 independent experi ments for each group. 2-way ANOVA, $F(2,28)=3.726$, $p=0.0367$. Multiple comparison for Cas9+ : empty vs gRNA1 p=0.7, empty vs gRNA4 p=0.0059, gRNA1 vs gRNA4 p=0.0320. Tukey's post-hoc test.
B. No significant changes in the levels of other miRNAs after Cas9 inactivation. In the same animals as in Fig 1C, we also measured the levels of miR-9-5p, miR-93-5p, miR-128-3p, miR-132-3p, miR-137-5p and miR218-5p. Expression of gRNA4 did not have any effect compared to empty vector in Cas9$^-$ or in Cas9$^+$ cells.

**Roura-Martinez et al Suppl Figure 5**

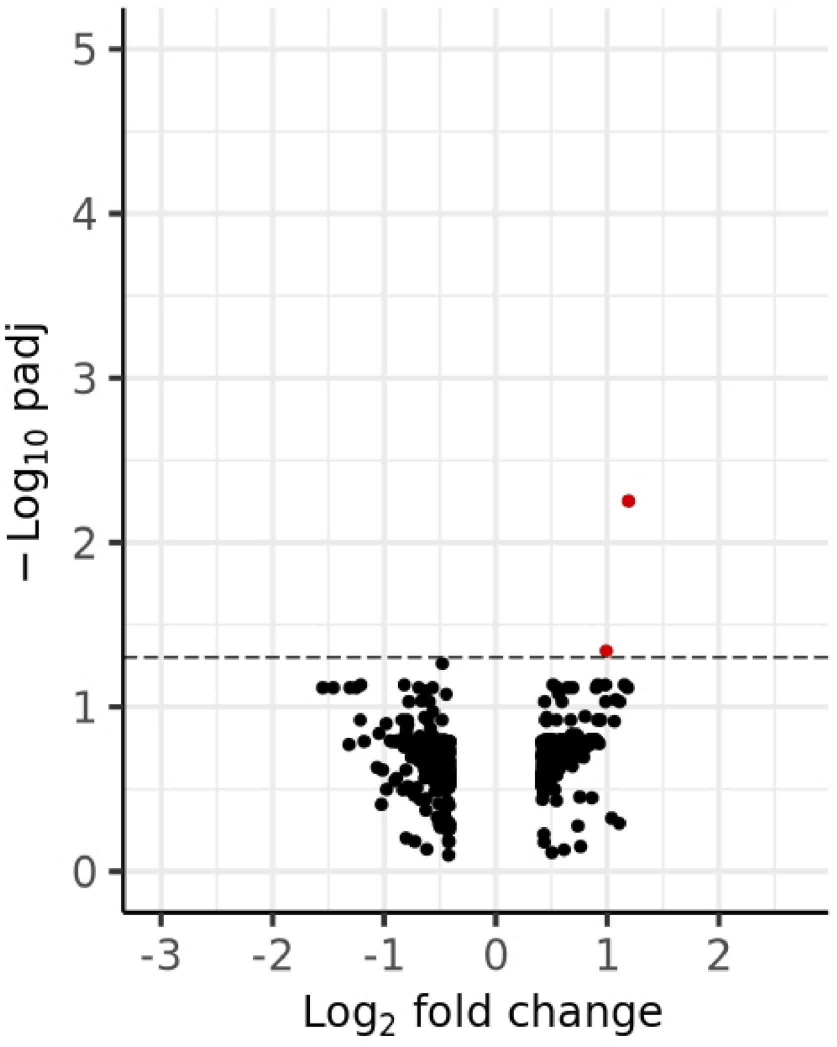


Supplementary Figure 5. Inactivation of miR-124 in vivo does not alter gene expression in uninfected Cas9- nuclei. Vol cano plot show almost no differentially expressed genes in Cas9- nuclei between animals injected with AAV-miR-124-Cas9$^{empty}$ and AAV-miR-124-Cas9$^{g4}$.
